# Isolation-by-distance and population-size history inferences from the coho salmon (*Oncorhynchus kisutch*) genome

**DOI:** 10.1101/2022.06.14.496192

**Authors:** Eric B. Rondeau, Kris A. Christensen, David R. Minkley, Jong S. Leong, Michelle T.T. Chan, Cody A. Despins, Anita Mueller, Dionne Sakhrani, Carlo A. Biagi, Quentin Rougemont, Eric Normandeau, Steven J.M. Jones, Robert H. Devlin, Ruth E. Withler, Terry D. Beacham, Kerry A. Naish, José M. Yáñez, Roberto Neira, Louis Bernatchez, William S. Davidson, Ben F. Koop

## Abstract

Coho salmon (*Oncorhynchus kisutch*) are a culturally and economically important species that return from multiyear ocean migrations to spawn in rivers that flow to the Northern Pacific Ocean. Southern stocks of coho salmon have significantly declined over the past quarter century, and unfortunately, conservation efforts have not reversed this trend. To assist in stock management and conservation efforts, we generated two chromosome-level genome assemblies and sequenced 24 RNA-seq libraries to better annotate the coho salmon genome assemblies. We also resequenced the genomes of 83 coho salmon across their North American range to identify nucleotide variants, characterize the broad effects of isolation-by-distance using a genome-wide association analysis approach, and understand the demographic histories of these salmon by modeling population size from genome-wide data. We observed that more than 13% of all SNPs were associated with latitude (before multiple test correction), likely an affect of isolation-by-distance. From demographic history modeling, we estimated that the SNP latitudinal gradient likely developed as recently as 8,000 years ago. In addition, we identified four genes each harboring multiple SNPs associated with latitude; all of these SNPs were also predicted to modify the function of the gene. Three of these genes have roles in cell junction maintenance and may be involved in osmoregulation. This signifies that ocean salinity may have been a factor influencing coho salmon recolonization after the last glaciation period – generating the current pattern of variation in these three genes.

## Introduction

Coho salmon have special cultural significance to the people of the First Nations in British Columbia and have traditionally been one of the highest-value Pacific salmon in the commercial and recreational fishery sectors. In 1977, a climatic regime shift in the North Pacific Ocean ushered in three decades of increasing global salmon production that culminated in 2009, when over 600 million salmon (1.1 million metric tonnes) were harvested [1]. However, this increased production of salmon masked substantial variability in regional abundances and species composition. Whereas the productivity and harvest of chum (*Oncorhynchus keta*), pink (*Oncorhynchus gorbuscha*), and sockeye salmon (*Oncorhynchus nerka*) increased throughout the North Pacific after 1977, the opposite was true for coho and Chinook salmon (*Oncorhynchus tshawytscha*). These declines became particularly acute after 1989 when marine survival for these species began a downward spiral that has yet to be reversed [1, 2]. A severe decline in the highly lucrative recreational coho salmon fishery in the Strait of Georgia saw the numbers of fish caught decline from an average of over 500,000 to less than 100,000 throughout the 1990s [3]. In 2004, the recreational catch in the Strait of Georgia was 9,500 coho salmon [4].

In British Columbia (BC), the Salmon Enhancement Program (SEP) was launched to double salmon production with the establishment of 18 major Department of Fisheries and Oceans (DFO) operated hatchery facilities and spawning channels. Throughout a stable harvest period of the 1980s, SEP releases of coho salmon juveniles increased from 5 to over 20 million, and the proportion of hatchery salmon in the fisheries increased from 5 to 20%. A precipitous decline in coho salmon production occurred in the 1990s. The decline was even more dramatic in the inner waters of the Strait of Georgia and Puget Sound. By the end of the decade, the commercial coho salmon fishery was closed and ‘marked or hatchery-only’ recreational fisheries had been instituted as a wild coho salmon conservation measure in southern BC.

The concern that hatchery fish were replacing wild fish was raised and, indeed, by 1998 70% of coho salmon in the Strait of Georgia were of hatchery origin [5]. From 1998 to 2007, the survival of the Strait of Georgia coho salmon over the first four months after entering a marine environment (May-September) decreased from 15% to 1% [2]. Processes associated with the low early marine survival remains unknown, but marine climatic changes were implicated and hatchery salmon survival was even lower than for wild salmon [2]. These results led to renewed calls for improved strategies for wild and hatchery coho salmon management, and a re-evaluation of wild-hatchery interactions in the species [2]. As such, a call was made to understand genetic influences on coho salmon survival and to produce high-quality genomic resources such as a chromosome-level reference genome assembly to enable technological support in informing management practices and decision within this species.

In a large-scale coho salmon population structure analyses of coho salmon sampled from 318 localities, in 38 different regional groups in North America and Russia (representing most of the natural distribution of coho salmon), 17 microsatellite loci showed that salmon clustered geographically and regions could be delineated along a north – south gradient, with reduced variation to the north and isolated inland populations [6]. These results were refined with increased genetic markers, finding that isolation-by-distance from a main southern glacial refugia after the last ice-age could explain most of the patterns of genetic diversity in modern coho salmon across the North American distribution [7, 8]. These last two studies were supported by the reference genome assemblies described in this study and illustrate how important such resources are for understanding the basic biology of a species.

With this in mind, the goals of this study were to expand upon our basic understanding of the coho salmon genome and help build upon the knowledge of the already excellent framework of population structure mentioned above. Our method to do this was to construct a high quality, annotated reference genome assembly and by building a comprehensive inventory of genetic variation (SNPs) from a wide geographical distribution. From the complete SNP dataset, we were then able to expand upon what was known about isolation-by-distance and demographic history of coho salmon. RNA-seq data for various tissues were also generated to facilitate genome annotation by the NCBI.

## Materials and Methods

### Coho salmon samples for genome assembly

All animals were reared in compliance with the Canadian Council on Animal Care Guidelines, under permit from the Fisheries and Oceans Canada Pacific Region Animal Care Committee (under Ex.7.1). Using Inch Creek coho salmon, we generated fully homozygous diploid gynogenetic individuals (doubled haploids) to help improve genome assembly quality (as noted in [9]). For details on doubled haploid generation and DNA extraction methods, please see the Supplemental Methods section. Tissues were also collected for RNA-seq and included kidney, heart, head kidney, spleen, gill, nares, ovary, white muscle, brain, eye, gut, liver, skin, stomach, and pyloric caecum. See the Supplemental Methods for further details on RNA extraction.

### Genome sequence and assembly – Version 1

A common sequencing and assembly pipeline for salmonids was used for this version of the genome assembly (e.g., [10–12]). Full details of sequencing and genome assembly can be found in the Supplemental Methods section. A brief description of the assembly involved generating Illumina libraries (mate-pair and paired end), generating PacBio data, and assembling the Illumina sequence data using Allpaths-LG [13] followed by scaffolding with PacBio data using PBJelly [14]. All sequencing data related to this genome assembly and annotation were submitted to the NCBI under BioProject Accession: PRJNA352719. Genome completeness was assessed using BUSCO (v3.0) [15], with default settings aside from “-sp zebrafish” using the ODB9 Actinopterygian database.

A circos plot (v0.69-4) was generated to show the relationship of homeologous chromosome resulting from the salmonid specific genome duplication [16, 17] for both versions of the genome assembly. For further details of the circos plot see Supplemental Methods.

### Genome sequence and assembly – Version 2

Version 2 of the coho salmon genome assembly incorporated 10X chromium data, Hi-C data, and new PacBio data with the previous Illumina sequencing and PacBio data generated for the first version. Some of this data came from a different doubled-haploid coho salmon individual compared to the first version. For full details of sequencing and assembly methodology see Supplemental Methods.

### Transciptome and annotation

RNA-seq data was generated from 15 tissues taken from the same doubled-haploid coho salmon used to produce the first genome assembly. RNA-seq data was also generated from two other coho salmon for this project, including: spleen, head kidney, kidney, gill, and gut (gut was only from one of the two salmon). In total, RNA from 24 tissues were sent for library construction and sequencing at the Michael Smith Genome Sciences Centre (Vancouver, BC, Canada). Eukaryotic single-strand RNAseq libraries were prepared and sequenced across 7 lanes of PE125 sequencing on an Illumina HiSeq2500. Four tissues were pooled per lane except brain, ovary, liver and gut from the genome individual (DH3), which were pooled two per lane. Sequences were submitted to NCBI under SRR5333359-SRR5333382 for eventual inclusion in the standard NCBI Eukaryotic Genome Annotation pipeline, which has been used on many genome assemblies. This data was generated for use in the NCBI annotation (Version 2), but was not used in any other way in this study.

### Repeat library

A species-specific repeat library was generated for coho salmon using the methodology developed for salmonids in [18], and fully described in [10]. In brief, the Atlantic salmon repeat library [18], was combined with repetitive sequences from the RepBase database [19]. The RepBase sequences were derived from the Salmoniformes family. They excluded simple repeats. RepeatModeler v1.0.8 [20] was also used together with the genome assembly in a *de novo* approach. The repetitive sequences were then aligned to the coho genome with BLASTN [21]. Sequences were classified into either high-confidence or low-confidence categories based on frequency and length. Low-confidence repeats were removed, and after filtering all of the sequences were compared to each other using an all-by-all BLASTN search. A redundancy filter was applied, prioritizing longest and highest-confidence repeats where two sequences were considered to overlap.

### Whole-genome resequencing and nucleotide variant calling

Whole-genome resequencing was used to characterize broad genomic characteristics across the coho salmon’s North American range. Table 1 contains a list of sampled locations (see File S1 for more information). We included one commercial strain from Chile as well (Table 1).

**Table 1.** Whole-genome resequencing sources

| Source | Country | State/Province | Count |
| --- | --- | --- | --- |
|  |  |  | Female, Male |
| Klamath River (Hatchery) | US | CA/OR | 1F, 4M |
| Deschutes River (Hatchery) | US | CA/OR | 2F, 3M |
| Big Quilcene River (Hatchery) | US | WA | 2F, 3M |
| Wallace River (Hatchery) | US | WA | 6M, 4? |
| Tsoo-Yess River (Hatchery) | US | WA | 1F, 4M |
| Inch Creek (Hatchery) | Canada | BC | 3F, 5M |
| Capilano River (Hatchery) | Canada | BC | 5F |
| Robertson Creek (Hatchery) | Canada | BC | 5M |
| Salmon River (Hatchery) | Canada | BC | 5F |
| Pallant Creek | Canada | BC | 5M |
| Kitimat River (Hatchery) | Canada | BC | 5F, 5M |
| Berners River | US | AK | 2F, 3M |
| Kwethluk River | US | AK | 1F, 4M |
| AquaChile (Strain) | Chile | NA | 5F |

State/Province Abbreviations: CA - California, OR - Oregon, WA - Washington, BC -
British Columbia, AK – Alaska

DNA was extracted from fin-clips using the DNeasy Blood and Tissue extraction kit (Qiagen) or a MagMAX DNA Multi-Sample Ultra Kit with a KingFisher (ThermoFisher Scientific). Following DNA extraction, samples were quantified by Qubit BR DNA assay (ThermoFisher) and integrity validated by agarose gel electrophoresis. At McGill University and Genome Québec Innovation Centre (Montreal, QC, Canada), individual Illumina libraries were constructed with Illumina TruSeq LT sample preparation kits, and each individual was sequenced separately on a lane of Illumina HiSeq2500 PE125 or in batches of four on a HiSeqXTen (PE150) lane, targeting approximately 15-30X coverage. Resequenced genomes were submitted to the NCBI under BioProject:PRJNA401427 and PRJNA808051 (File S1).

Nucleotide variant calling on the dataset followed GATK3 best practices where possible. BWA-MEM v0.7.17 [22] was used to align Illumina data to the reference genome (version 2), with -M option for Picard compatibility. The Picard v2.18.9 [23] AddOrReplaceReadGroups program was used to add read group IDs, and the MarkDuplicates program was used to mark duplicates (default settings). GATK v3.8 [24, 25] was then used to call genotypes. Base and variant recalibration were each performed once (for two rounds through genotyper). The variants used for recalibration were from 1) a reduced set of very high-confidence calls following default “hard-filtering” guidelines from GATK documentation from the first round of genotyping with a particular focus on coding regions, and 2) validated SNPs on a 200K Affymetrix SNParray [26].

Following genotyping, VCFtools v0.1.15 [27] was used to additionally thin data to only include biallelic SNPs with a minor allele frequency of 0.05 or greater, variants with fewer than 10% missing genotypes, and variants with a mean coverage between 5 and 200. Minor allele frequency was not used for filtering in the SMC++ analysis (below). Some individuals were removed at this point from the VCF file because they were not intended for this study (see NCBI BioProject: PRJNA808051 and File S1 for removed samples). They were included as it was more computationally efficient to call all individuals at the same time. Finally, the SNPs were filtered for linkage disequilibrium to reduce the influence of large haploblocks in the principal components analysis (PCA) (bcftools [28] version 1.9-102-g958180e +prune -w 20kb -l 0.4 -n 2).

### Whole-genome analyses

A PCA was performed with variants that had been filtered for linkage disequilibrium (see previous paragraph) using PLINK [29, 30] v1.90b6.15 with default parameters. PLINK was also used to identify and quantify runs of homozygosity using default settings (Figure S1). For comparison, we also performed the analysis with the following parameters (Figure S2): min SNP count – 100, min length – 100 kb, max inverse density – 50 kb/SNP, max internal gap 100 kb, max heterozygous genotypes 1, SNP scanning window size – 100, min scanning window hit rate – 0.05, max missing calls – 20.

Private allele counts per river were tallied using the populations module in Stacks [31, 32] version 2.54 with default parameters. Populations with more than five individuals were randomly subsampled to five to reduce the influence of uneven sampling on the number of private alleles identified. Stacks was also used to calculate other population level metrics such as observed heterozygosity, nucleotide diversity (Pi), and Fis with default settings.

A genome-wide association (GWA) analysis was performed to characterize the extent of isolation-by-distance previously reported by several authors (e.g., [7, 33]). The trait of interest under investigation was latitude (the Chile strain of coho salmon was excluded from this analysis). We used PLINK with default settings to perform this analysis. Population structure was not included in this analysis as a covariate because we were trying to characterize the fraction of the genome with a north – south gradient and adding this covariate would remove much of that variation. R [34] and the qqman package in R [35] were used to visualize the GWA analysis.

We tested for gene ontology enrichment based on the annotated variants that were associated with the north – south gradient (for variants that were ‘moderately’ likely to influence gene function and for those having ‘low’ or ‘moderate’ likelihood). SnpEff [36] version 5.0e and the gene annotation from the NCBI were used to annotate nucleotide variants for potential function alterations using default settings. Blast2GO [37] and OmicsBox [38] version 2.0.36 were used to test for enriched GO categories using default parameters.

To infer demographic histories of the salmon from the various rivers, we used SMC++ [39] version 1.15.4.dev18+gca077da. In this analysis, we set the mutation rate to 8e-9 bp/generation and the generation time to 3 years. These parameters were previously used in another coho salmon study examining demographic histories [7]. We used nucleotide variants that were not filtered for rare variants (e.g., MAF < 0.05). We also used the --missing-cutoff option (50 kbp) in SMC++ to reduce the influence of missing genotypes (e.g., in centromeres).

## Results

### Genome assemblies

The size of both versions of the coho salmon genome assembly was 2.3 Gb, which is also the same size of the closely related Chinook salmon (*O. tshawytscha*) genome assembly [40]. However, version 2 of the coho salmon genome assembly was much more contiguous than version 1 and had a more complete gene set (inferred from BUSCO completeness). There was an almost 20x fold increase in contig N50 between version 1 (58 kb) and version 2 (1,159 kb) of the genome assembly (Table 2). Likely as a consequence of the increase in contiguity, the number of complete BUSCOs rose from 91% to 99%, which is comparable to the human genome assembly at 99% [41]. The proportion of repeats also rose from 44.82% to 53.12% (compared to 52.94% in Chinook salmon), and the number of annotated genes increased from 41,179 to 60,330 (47,105 in Chinook salmon). The NCBI reported that from version 1 to version 2, 37% of the genome annotations were new and that 16% of the annotations on version 2 required major changes from the previous version [42]. We note that the genome assembly was produced from sequence data from two coho salmon and therefore not haplotype resolved but chimeric in nature.

**Table 2.** Genome statistics

|  | Contig N50 | Contig # | BUSCO | % Repeats | Genes |
| --- | --- | --- | --- | --- | --- |
| Ver 1 | 58,118 | 97,074 | 91%-55:37* | 44.82† | 41,179† |
| Ver 2 | 1,159,298 | 8,770 | 99.2%-57.1:42.2*† | 53.12† | 60,330† |

Ver 1, NCBI: GCF\_002021735.1; Ver 2, NCBI: GCF\_002021735.2
\*Percent complete-single:duplicate
†Reported by NCBI (NCBI used actinopterygii\_odb10 for BUSCO)

The coho salmon genome has extensive signatures of chromosomal duplication (Figure 1, Table 2), which have been retained from the whole genome duplication common to all salmonids [17]. The majority of duplicated regions from the salmonid-specific genome duplication have diverged to a point where it is relatively easy to differentiate between the copies (Figure 1, ≤90% identity), but certain sections of the genome have retained high-sequence similarity where it is difficult to distinguish between copies (Figure 1). Regions with very high-sequence similarity remain as unplaced scaffolds as it was not possible to resolve which sequence belonged to which duplicated region (see assembly methods; available on the NCBI website [43]). The number of duplicate BUSCOs increased from 37% to 42.2% between versions (Table 2), which suggests that the second assembly was able to distinguish between similar paralogs/homeologs better whereas the first assembly likely collapsed them into a single gene/BUSCO.

**Figure 1.**
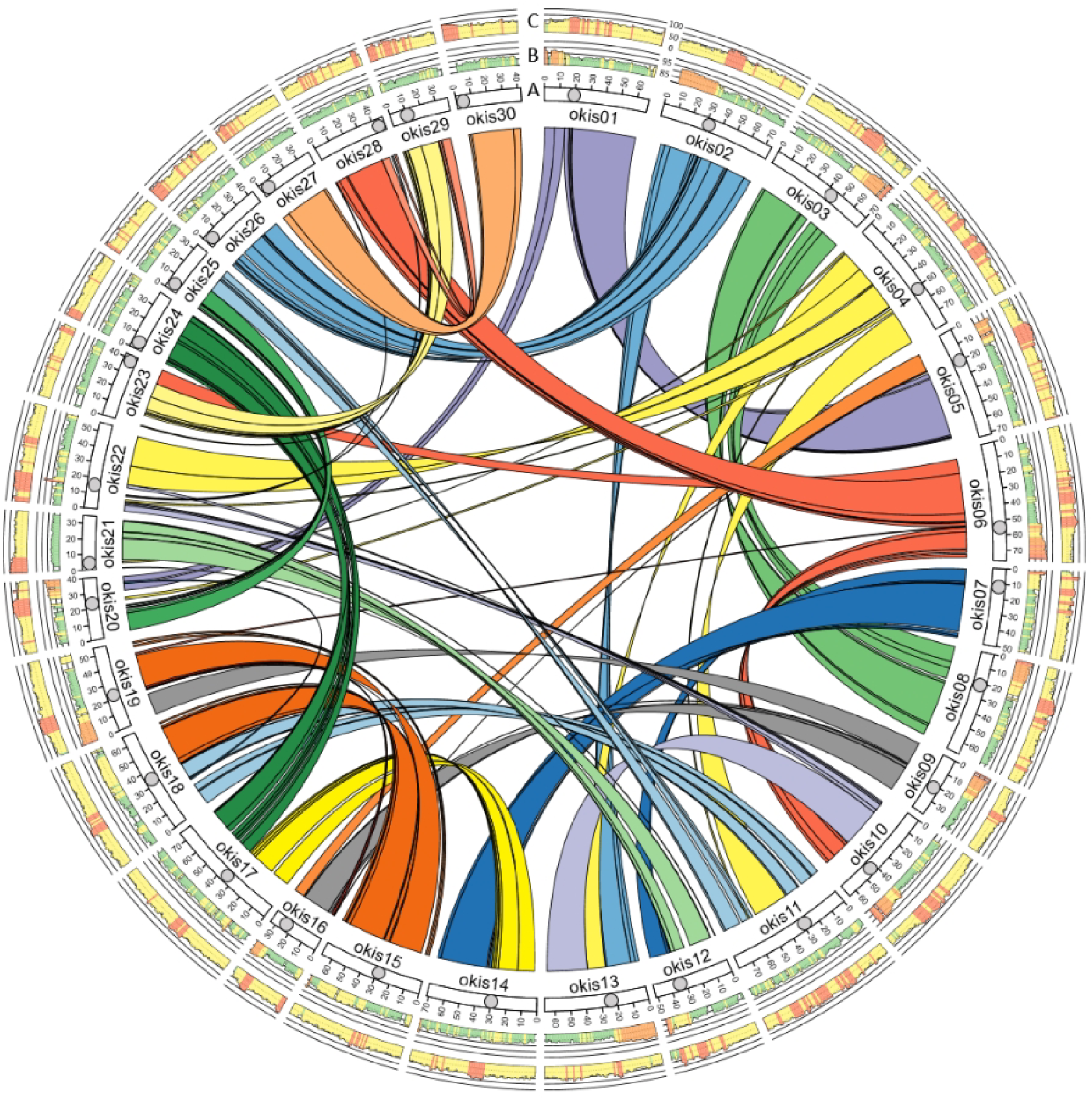
Circos plot of the first version of the coho salmon genome assembly. In the interior of the Circos plot are the links between duplicated regions of the chromosomes/linkage groups (i.e., homeologous regions). A) Representations of the chromosomes with the approximate position of the centromere marked by a filled circle. The tick marks represent 10 Mbp intervals. B) The percent identity between duplicated regions of the chromosome. The orange-red color represents very high similarity (> 94%), the orange color high similarity (91-94%), the yellow moderate (88-91%), and the green low (< 88%). C) The fraction of repetitive elements, with red representing high (> 60%), yellow as moderate (35-60%), and green as low (< 35%).

The coho salmon genome also has a high retention of repetitive elements (Figure 1, Table 2), which is another commonality of studied salmonids (e.g., [12, 18]). This is especially true in regions near the centromere where the fraction of repetitive elements is roughly 75% (Figure 1). That value is high compared to the genome average of 53% (Table 2). For comparison, the most recent version of the Chinook salmon genome also has a repeat content of 53% [44].

### Population genomics

A PCA of 83 resequenced coho salmon genomes sampled from across North America (and aquaculture samples), revealed that coho salmon clustered by region with the exceptions of the Salmon River and Inch Creek (Figure 2). On the first principal component of the PCA, the Salmon River clustered away from all the other samples. This river belongs to the Thompson River watershed, and coho salmon from this region have previously been observed to cluster in a similar manner [7]. Inch Creek salmon might cluster separately as an artifact since the genome assembly was derived from an Inch Creek salmon. This might increase read-alignment scores and influence SNP-calling in some regions of the genome.

**Figure 2.**
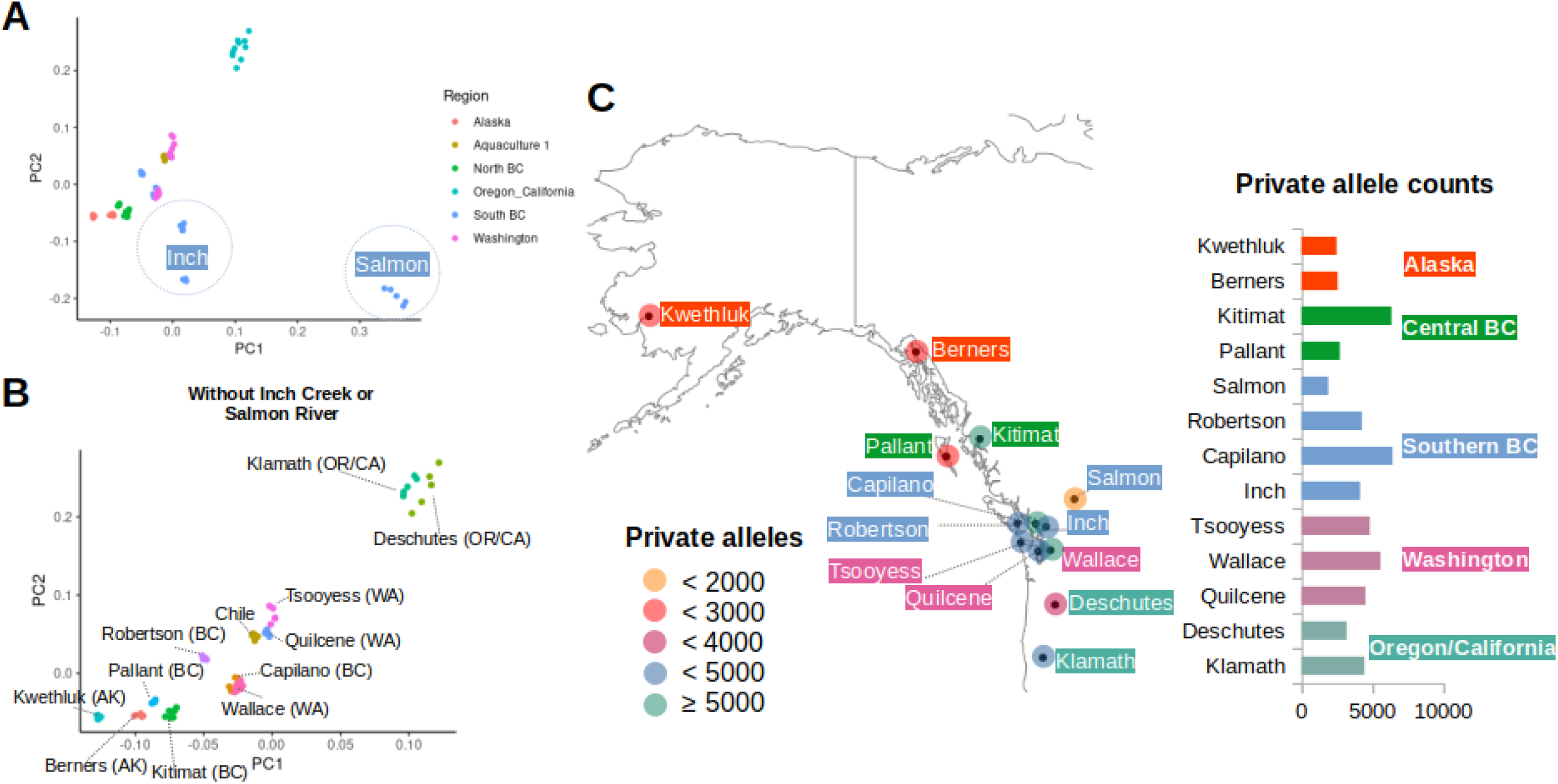
Influence of isolation-by-distance on coho salmon population structure. A) A PCA of coho salmon based on variants that were filtered for linkage disequilibrium (plotted using ggplot [69]). Circles were drawn around Inch Creek and Salmon River individuals to highlight that much of the variation of this PCA was due to differences of salmon from these rivers. B) The same figure as A, with Salmon and Inch Creek salmon removed. When Inch Creek and Salmon River salmon are removed from the graph, the influence of latitude/isolation-by-distance can be observed on PC1 and PC2. Individuals from the same river (same colour) also tended to cluster near each other. C) A map of the various rivers sampled for this study and the corresponding private allele counts (plotted with the maps [70] package in R). The private allele counts are also displayed to the side as a bar graph.

Excluding the Salmon River and Inch Creek samples, all other samples clustered by region and by latitude in a manner consistent with isolation-by-distance suggested by [7]. The Salmon River group have the lowest private allele counts (1,876 vs. a median of 4,188) and observed heterozygosity (0.22966 vs. a median of 0.285565). They also have the highest total runs of homozygosity (Figures S1 and S2). The region with the highest private allele count appears to be around the Puget Sound (e.g., Wallace River, private allele count = 5,546) and Strait of Georgia regions (e.g., Capilano River, private allele count = 6,415). Most of the northern rivers have low private allele counts with the exception of the Kitimat River (private allele count = 6,341), which has the second highest count (Figure 2).

To investigate how much of the genome has been influenced by isolation-by-distance, we quantified the number of SNPs associated with the latitude gradient observed from the PCA above (Figure 3, File S2). Roughly 13.9-33.8% of the 5,631,459 variants were associated with latitude at a significance level of 0.01-0.1 without multiple test corrections (Figure 3). The proportion of variants associated with latitude dropped to 0.07% after the alpha threshold was set to 0.05 with a Bonferroni correction (these variants were widely distributed throughout the genome).

**Figure 3.**
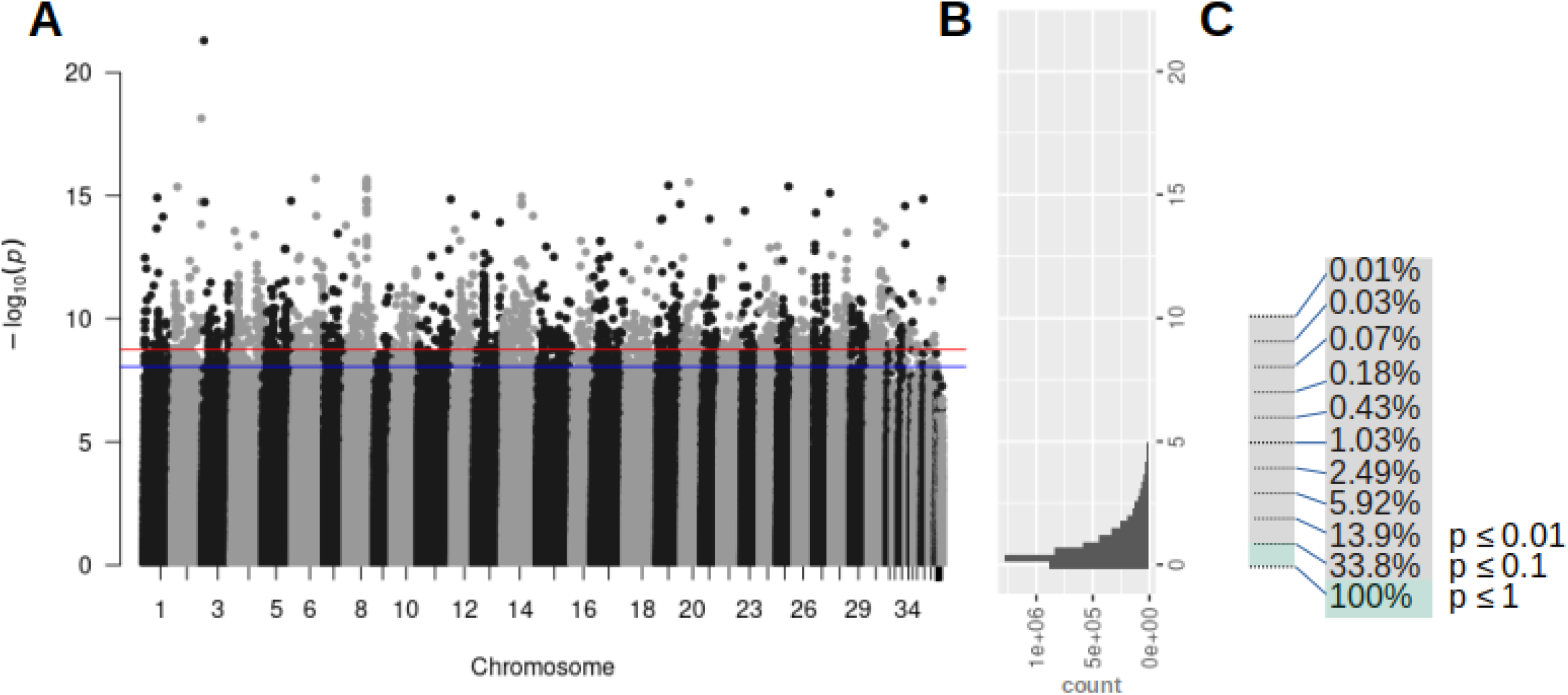
Fraction of the genome responsive to isolation-by-distance. A) A Manhattan plot of the latitude genome-wide association analysis of all coho salmon except for the Chile strain. The red line represents a significance level of 0.01 after a Bonferroni correction and the blue line a 0.05 level with the same correction. B) A histogram of variant counts for different significance levels. C) The percent of the variants at different significance levels. The percentage represents all the variants at or above the line. There are 33.8% of the variants that have a significant association with latitude at *p* ≤ 0.1 without multiple test correction.

In Table 3, the most common nucleotide variant annotations from SNPeff are shown, with intronic and intergenic variants being the most common type of variant annotation. The variants that were significantly associated with latitude (see previous paragraph, 0.07%) have a similar broad distribution of annotations relative to the entire genome rather than enriched for variants that are likely to influence gene function (Table 3, File S2). For instance, the percent of intergenic nucleotide variants remained at 31.2% of the total number of variants for the whole genome and for variants that were significantly associated with latitude (Table 3). We would expect that if variants were influencing traits under selection (e.g., based on latitude), the distribution would change between all variants and those significantly associated with latitude if those SNPs influenced gene function (e.g., 3’ UTR and missense annotations).

**Table 3.** Distribution of common nucleotide variant annotations

|  | Entire genome* | Associated with<br>latitude* |
| --- | --- | --- |
| Intron | 44.0% | 42.1% |
| Intergenic | 31.2% | 31.2% |
| Upstream | 10.4% | 11.2% |
| Downstream | 7.3% | 7.6% |
| 3' UTR | 2.8% | 3.2% |
| Missense | 0.8% | 1.7% |
| Synonymous | 1.1% | 1.1% |

\*5,631,459 genomic SNPs, 3,940 significant SNPs from GWA analysis

Significant latitude-associated nucleotide variants (0.07%) identified in the GWA analysis that were annotated as having a ‘Moderate’ likelihood to influence gene function by SnpEff were found in 45 genes (File S2). Of these 45 genes, 4 genes had two or more variants that were annotated as ‘Moderate’ in their impact on gene function (Figure 4). No enriched gene ontologies were identified from genes with ‘Moderate’ or even ‘Moderate’ + ‘Low’ (87 genes) nucleotide variant annotations (File S2). Only when all genes with associated variants were tested, regardless of influencing function, did we observe enriched GO terms (data not shown).

**Figure 4.**
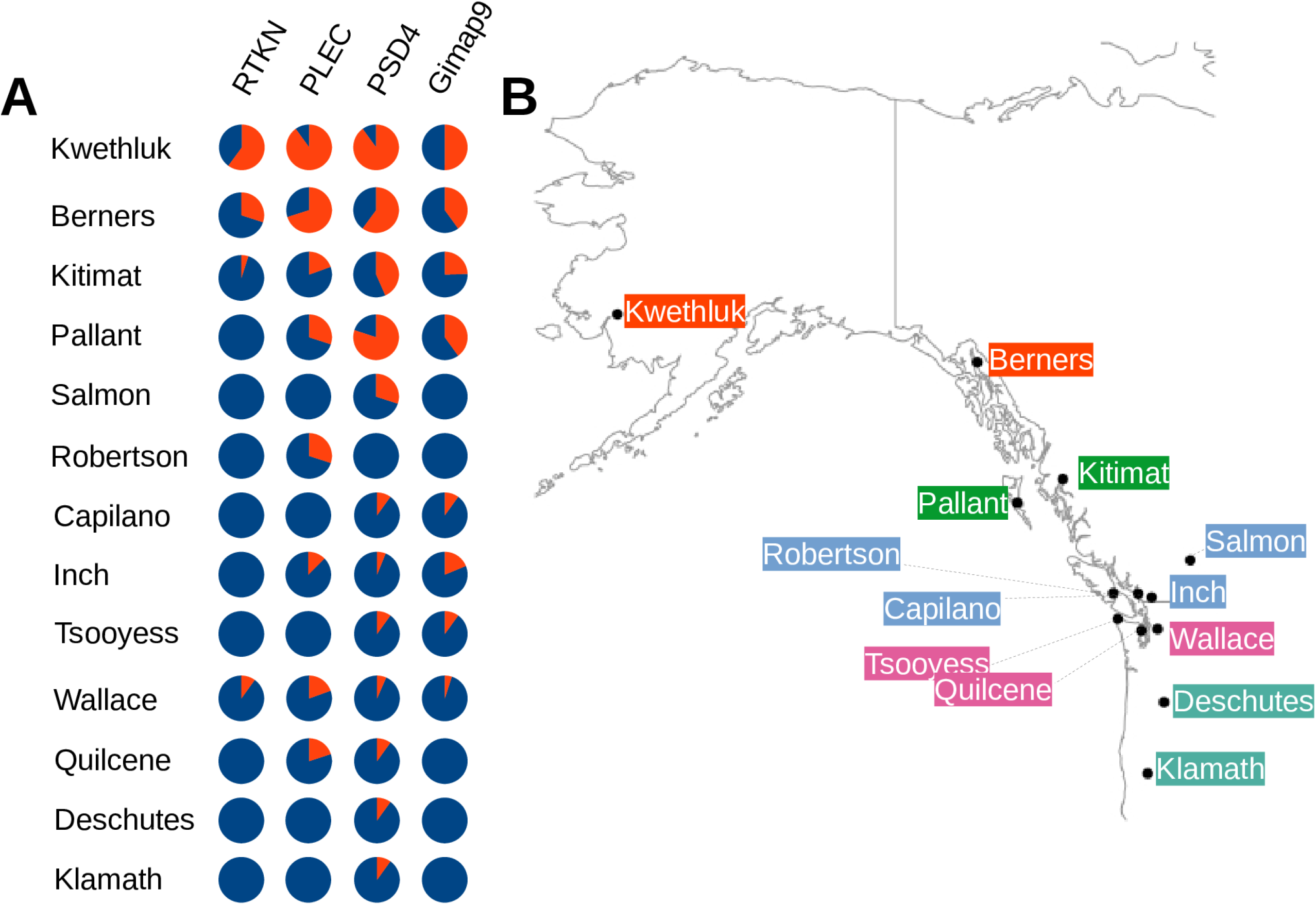
Genes with multiple ‘moderate’ nucleotide variants that may influence function. Four genes were found to be associated with latitude and also contained multiple SNPs that likely modify gene function. The four genes are: rhotekin-like (RTKN LOC109895613), plectin-like (PLEC LOC109904478), PH and SEC7 domain-containing protein 4-like (PSD4 LOC109868337), and GTPase IMAP family member 9-like (Gimap9 LOC109880231). A) Pie diagrams showing the distribution of reference (blue) and alternative alleles (red) for each gene and location. B) Map produced with the maps package in R showing the sampling sites.

To put the nucleotide variation generated by isolation-by-distance into a broader context, we identified possible times when northern populations could have recolonized after the last glaciation period. By modeling demographic histories from genome sequences using the SMC++ program, we were able to identify major decreases in effective population size (Ne) that correspond with the Cordilleran Ice Sheet maximum and the presumed penultimate global glacial maximum (Figure 5). We also observed that for some populations, mostly northern, there was an additional drop in effective population size between 3,750 and 8,000 years ago (Figure 5).

**Figure 5.**
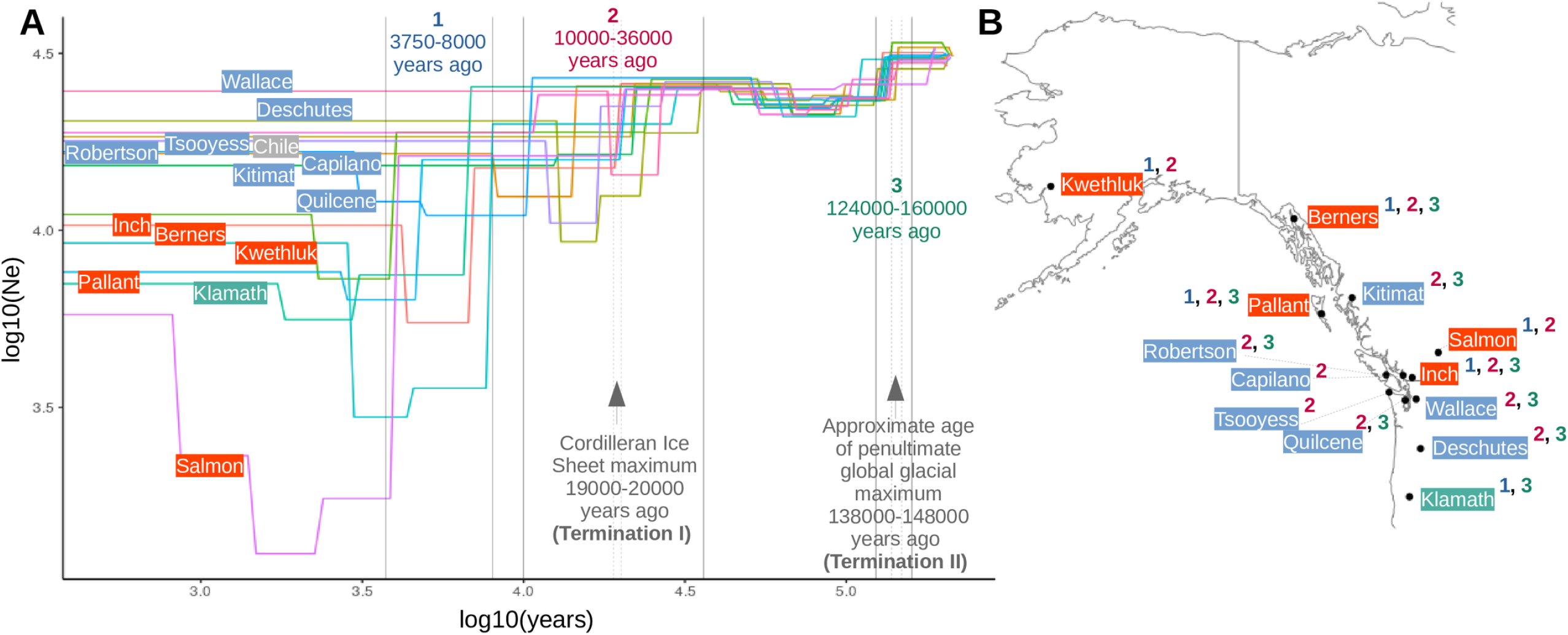
Demographic histories of coho salmon populations based on genome sequencing. A) Each labeled line represents multiple individuals from the same river or strain. The X-axis represents calendar years based on a generation time of 3 years for coho salmon. The Y-axis is the effective population size (Ne) estimate. The estimated age of the Cordilleran ice sheet maximum was taken from [50]. The approximate age of the last interglacial period was based on [71]. B) For each location, a number nearby indicates a drop of at least 5,000 in the Ne for one of the time points noted in A. The colour of the river label indicates if the river had a drop in Ne during the 1^st^ time interval (orange – yes, blue – no, the Klamath River was the only southern river with a drop during the 1^st^ time period).

## Discussion

As with previous analyses of salmonid genomes [10, 12, 17, 18, 45, 46], the retention of duplicated chromosomes (i.e., homeologs) from the salmonid-specific whole genome duplication [17] is a defining feature of the coho salmon genome. Some of the duplicated regions have likely retained very high sequence similarity for roughly 90 million years (time estimate from [17, 45, 47]). A possible mechanism for high sequence similarity retention is through tetrasomic inheritance [48].

The second version of the coho salmon genome assembly resolved a greater number of duplicated regions of the genome compared to the first version. The better resolution of duplicated regions can be observed with the increase in gene count and the number of duplicated BUSCOs identified. Finer detail in these regions may help us in future studies to better understand the residual impacts of whole genome duplication on the biology of salmon.

From resequenced coho salmon genomes, we were able to better understand population structure of coho salmon and its relationship with isolation-by-distance. One of the striking features of the PCA of coho salmon populations was how divergent Salmon River salmon were to all other populations. The Salmon River is part of the Thompson River watershed and coho salmon from this system were thought to be isolated from all other populations for potentially 150,000 years before secondary contact roughly 13,500 years ago (essentially during the previous glacial period) [7]. This would be consistent with findings in kokanee (*O. nerka*, a landlocked sockeye salmon ecotype) in the upper Columbia River that similarly appear divergent from all other populations of sockeye salmon and kokanee [12]. Taken together, these pieces of evidence might be interpreted as support for a glacial refugium near the intersection of the Cordilleran Ice Sheet and the Laurentide Ice Sheet.

A more likely alternative is that another unknown factor was influencing past analyses and the PCA from the current study. The Salmon River coho salmon have increased runs of homozygosity, reduced heterozygosity, and reduced private alleles, which are indicators of a recent and extensive bottleneck. We were also able to infer the demographic history from whole genome sequences of the Salmon River coho salmon and found evidence of a bottleneck (from ^~^Ne 16,227 to ^~^Ne 1,749) around 4,000 years ago. These results could help explain why the Salmon River coho salmon appear so divergent in a PCA, as low genetic diversity might be expected to increase the amount of variation in the analysis since most of the other individuals do not have low genetic diversity. We only collected samples from one tributary of the Thompson River (a part of a much larger basin) and can only suggest that a plausible hypothesis from this data is that recolonization of the Salmon River from a small founding population took place after glaciers receded. We did not account for the influence of hatcheries, which could also influence many of the metrics discussed above. Also, we did not incorporate linked selection in demographic modeling as the type of analysis that we used was not amenable. Without linked selection accounted for, there could be biases in times and effective population sizes from our estimates [49]. Based on all the genetic diversity metrics (above), demographic modeling, and what has previously been published on the time of the most recent glacial maximum [50], however, recent recolonization of the Salmon River remains a likely alternative hypothesis to a glacial refugium between ice sheets.

Most other streams, except Inch Creek, clustered in the PCA based largely on latitude for both PC1 and PC2 of the PCA. With a much more extensive sampling strategy, Rougemont et al. (2020) found a similar trend and tested various demographic histories [7]. The authors of that study found that the best supported model was a glacial refugia to the south with recolonization of the northern streams after glacial retreat – generating genomic signatures of isolation-by-distance. The private allele analyses from the current study also supports this interpretation. The private allele analysis identified that most of the northern streams have low private allele counts compared to southern streams.

To better understand the pattern of genomic isolation-by-distance along the latitude gradient, we performed a GWA analysis based on stream latitude. We found that 13.9% of the variants were associated with latitude based on a *p*-value of 0.01 without multiple test correction. To put this into perspective, the north – south gradient likely formed after the last Cordilleran Ice Sheet maximum 19-20,000 years ago [50] but could have formed later between 3,750-8,000 years ago based on our demographic history modeling. This would suggest that a large fraction of the nucleotide variants responded within less than 8,000 years to the influence of isolation-by-distance.

We analyzed the influence of significantly associated nucleotide variants on gene function and also the distribution of annotated variants to better understand if selection played a large role in establishing the north – south gradient. We tested if genes with significantly associated variants (α = 0.05, Bonferroni-correction), with ‘Low’ to ‘Moderate’ likelihoods of influencing gene function, belonged to any enriched GO categories. If a trait was under selection based on latitude, we might expect enriched GO terms associated with that trait. We did not find any enriched GO categories from the GWA analysis. This may indicate that selection may not have contributed much to establishing the north – south gradient.

When comparing the distribution of the most common variant annotations of the full dataset with the variants that were significantly associated with latitude, the largest fold difference was between the missense mutations (Table 3). We observed a ^~^2x difference from 0.8% missense mutation rate in the entire genome to 1.7% in the variants associated with latitude. While this increase does suggest selection may have contributed to the development of the latitude gradient, we interpret that, because the majority of the other annotations have similar frequencies, the majority of the variants that make up the gradient are not under direct selection. Further, the increase in missense mutations may represent slightly deleterious variants that escaped selective pressure during postglacial recolonization and expansion. Linked selection could still play a larger role but was not investigated here.

With or without selection, the north – south gradient of nucleotide variants likely influences some phenotypic differences in a similar gradient. As an example, we identified genes that had multiple nucleotide variants that are predicted to moderately influence gene function, and which also have an association with latitude. These included the rhotekin-like (*RTKN*, unknown function [51]), plectin-like (*PLEC*, giant cytoskeleton scaffold [52]), PH and SEC7 domain-containing protein 4-like (*PSD4*, tight junctions maintenance [53]), and GTPase IMAP family member 9-like (*Gimap9*, possibly involved in T-cell development [54, 55]) genes. The nucleotide diversity of these genes largely arises from the frequency of the alternative allele in the four most northern streams – regions that would have likely been recolonized most recently assuming a main southern glacial refugium.

Interestingly, *PLEC* is a candidate gene associated with migration distance in brown trout (*Salmo trutta*), perhaps through its role in osmoregulation [56]. Cells without *PLEC* were found to be more sensitive to changes in osmolarity (shrinking more after exposure to urea) [57], and hatch-stage whitefish (*Coregonus lavaretus*) exposed to high salinity have significantly higher *PLEC* protein expression [58]. Two of the other genes with multiple variants moderately-likely to modify gene function and which were associated with the north – south latitudinal gradient, *RTKN* and *PSD4*, may also have roles in salinity tolerance. An Atlantic cod (*Gadus morhua* L.) nucleotide variant in the intron of *PSD4* was found to be associated with a salinity gradient between the North Sea and the Baltic Sea [59]. Likewise, researchers discovered that Rho (RTKN is an effector protein of RhoA [51, 60]), is activated by hyperosmotic stress [61].

*PLEC, PSD4*, and *RTKN* all appear to be involved in cell junction functionality. Cell junctions observed in a *PLEC* knockout cell line appeared to be compromised [62], *PSD4* (also known as EFA6B) is required for efficient tight junction formation [63], and *RTKN* influences cell junctions through PIST [51] and Septin proteins [51, 64]. It is thought that cellular tight junctions play an important role in water and salt balance in teleost fishes [65]. Considering that three of the four genes with multiple latitude associated nucleotide variants and which are moderately likely to alter gene function could impact cell junctions and osmoregulation, we hypothesize that ocean salinity may have influenced coho salmon recolonization of northern streams.

While, the Pacific Ocean salinity was thought to be ^~^4% higher during the last glacial maximum as freshwater was stored in glaciers [66], it is difficult to predict how the salinity gradient observed in modern times [67] might have been influenced as glaciers began to retreat (when northern recolonization would have been possible). If there was a difference in salinity between northern and southern regions, nucleotide variation in these genes may have facilitated northern colonization in some way.

From inferred demographic histories, we were able to estimate a recolonization date of some northern streams (based on the founder effect that would be expected to accompany recolonization) to between 3,750-8,000 years ago. This places an upper limit on the age of the latitude gradient and how swiftly such a gradient can form. These values are based on assumptions of a mutation rate of 8e-9 bp/generation and a generation time of three years. Linked selection may also bias our time and effective population estimates [49] as we did not account for them in modeling.

While it is important to remember that time and population estimates are influenced by many factors when inferring demographic histories from sequence data, multiple lines of evidence can be used to strengthen these inferences or put them in a more realistic context. Radiometric evidence supports that the Cordilleran Ice Sheet maximum occurred between 19,000 and 20,000 years ago [50]. Likewise, chemical properties of gases in Antarctic ice cores support this termination of the last glaciation period (Termination I) to roughly the same time period, as well as a previous termination of the penultimate glaciation period around 138,000 and 148,000 years ago (Termination II) [68]. In the demographic histories of the coho salmon, we noted dramatic declines of nearly all salmon populations for both these time periods. This observation supports the parameters used for modeling the demographic histories as we expect that populations might decline in response to increased glaciation or rapid climate change.

The overall trend we observed from modeling demographic histories was major drops in effective population size at each transition from glaciation to inter-glaciation period with increases for nearly all populations after the penultimate glaciation period and uncommon increases for specific rivers after the most recent glaciation period. At a species level, these transitional drops likely influence multiple aspects of coho salmon biology since genetic variability can contribute to many characteristics of a species.

## Conclusions

In this study, we generated two reference genome assemblies as tools for conservation and management of coho salmon. Additionally, we resequenced the genomes of a wide distribution of coho salmon from rivers along North America. We were able to identify a north – south gradient in the nucleotide variation of the genomes, which had been observed in previous studies. To add to previous observations, we quantified that approximately 13.9% of the variation in the resequenced genomes followed the north – south gradient. We also were able to estimate that the age of the north – south gradient is likely under 8,000 years of age. This gradient likely contributes to phenotypic diversity between northern and southern rivers since we identified gene modifying variants that were associated with the latitude gradient. Finally, we modeled demographic histories of the coho salmon from different rivers and discovered that major drops in effective population size were related to changes between glacial and inter-glacial periods. We believe the coho salmon genome assemblies will facilitate research to better understand coho salmon biology and may enhance management of this culturally and economically important species.

## Data availability

Raw data for the genome assembly was submitted to the NCBI under the BioProject PRJNA352719. Whole genome resequencing data was submitted under PRJNA401427 and PRJNA808051 to the NCBI BioProjects (see File S1 for specific samples used in this study). The VCF file used for analyses in this study was submitted to the GSA Figshare portal.

## Acknowledgements

We would like to acknowledge the extensive help from many individuals who contributed to work in sampling and sequencing in this work. For sequencing, we thank the staff at the McGill University and Genome Québec innovation Center as well as the Genome Sciences Centre who provided library prep and sequencing for PacBio long read libraries and individual coho resequencing work. We also thank the staff at the Michael Smith Genome Sciences Centre in Vancouver BC who performed library prep and sequencing for Illumina libraries used in genome assembly and RNA-seq libraries used in Genome annotation. The annotation work itself relied on the NCBI Eukaryotic Genome annotation pipeline and the wonderful staff who support this work. Sample collection involved a number of people, many of which remain anonymous to the authors, but we thank them for all their efforts. Coordinating sample collections were Heather Hoyt of the Alaskan Department of Fish and Game for samples from Alaska, Christian Smith at the US Fish and Wildlife’s Abernathy Fish technology Center for samples in Washington State and Oregon, John Carlos Garza at the NOAA Southwest Marine Fisheries Sciences Centre for samples collected in Klamath River, Justin Henry and Bruce Swift for samples from Aquaculture broodstock from Northern Divine Aquafarms Ltd (formerly Target Marine), and from Riverence LLC for Aquaculture broodstock. We would finally like to thank Compute Canada (Cedar) for computational resources.

## Author Contributions

EBR, KAC, JSL, and BFK performed genome assembly, chromosome assembly, genome submission, and generated genome metrics. DRM performed repeat library construction. EBR, CAD, MTC, AM, and DS performed wet-lab work including DNA and RNA extractions and mitochondrial sequencing. EBR, QR, EN, DRM, and JSL performed SNP calling and population genomic analyses. RHD, MTC, DS, and CB generated, raised, and dissected doubled-haploid samples for the genome assembly and transcriptome. REB, TDB, KAN, and JMY provided samples used in resequencing work. KAN provided early access to linkage map and additional guidance on its use. RHD, REW, TDB, KAN, JMY, RN, LB, WSD, SJMJ, and BFK initiated, planned and supervised the project. EBR, KAC, DRM and BFK wrote first draft of the manuscript.

## Supplemental Material

**Figure S1. Runs of homozygosity and admixture among coho salmon from different streams.** A) The top figure shows the total runs of homozygosity (ROH) for each individual (default settings in PLINK). The bottom figures show the admixture of each individual based on cluster counts of k=2 and k=3 (with Admixture software [72] using default settings with LD filtered SNPs). Streams are shown at the bottom and delineated by the alternating blue bar. B) A map (generated using the maps package in R) showing the locations in A.

**Figure S2. The relationship between runs of homozygosity and latitude.** A) Counts of runs of homozygosity (ROH) and the total length of the ROH when combined (see Methods for parameters used). There is a distinct cluster of Salmon River individuals with higher counts and lengths of ROH. B) The relationship between the average length of ROH per individual and latitude. The line was plotted using the geom_smooth function in ggplot2 with the linear model method. Latitude significantly (*p* = 0.029) explained variation in the average length of ROH (~6% of the variation, Adjusted R^2^ = 0.04886).

**File S1. Sample information and SRA accession numbers.**

**File S2. Nucleotide variants significantly associated with latitude.** The SignificantVariants tab in this spreadsheet file has information on all of the significantly associated SNPs with latitude. The ModerateGenes tab has information on SNPs that were both associated with latitude and also have annotations from SNPeff that were moderately likely to influence gene function. The Moderate+LowGenes tab has information on SNPs that were both associated with latitude and also have an annotation from SNPeff that were likely to have moderate or low influences on gene function. The GO tab has two lists of genes that were used in the GO enrichment analyses. The Frequency of MultiVariantGenes tab has information on the genes with multiple SNPs thought to moderately influence gene function and which were also associated with latitude. This information was used to generate pie charts. The GenotypeGenes tab has the genotypes for each individual for genes in the Frequency of MultiVariantGenes tab. The Regions tab has information on the latitudes used for each stream. The DistributionOfVariantAnnotations tab has information on SNPeff annotations from the SNPs that were significantly associated with latitude as well as the SNPs from the entire genome.

